# Temporal changes in plasma membrane lipid content induce endocytosis to regulate developmental epithelial-to-mesenchymal transition

**DOI:** 10.1101/2020.10.18.344523

**Authors:** Michael L. Piacentino, Erica J. Hutchins, Cecelia J. Andrews, Marianne E. Bronner

## Abstract

Epithelial-to-mesenchymal transition (EMT) is a dramatic change in cellular physiology during development and metastasis which involves coordination between cell signaling, adhesion, and membrane protrusions. These processes all involve dynamic changes in the plasma membrane, yet how membrane lipid content regulates membrane function during developmental EMT remains incompletely understood. By screening for differential expression of lipid-modifying genes over the course of EMT in avian neural crest, we have identified the ceramide-producing enzyme neutral sphingomyelinase 2 (nSMase2) as a critical regulator of a developmental EMT. nSMase2 expression begins at the onset of EMT, and *in vivo* knockdown experiments demonstrate that nSMase2 is necessary for neural crest migration. Further, we find that nSMase2 promotes Wnt and BMP signaling, and is required to activate the mesenchymal gene expression program. Mechanistically, we show that nSMase2 is sufficient to induce endocytosis, and that inhibition of endocytosis mimics nSMase2 knockdown. Our results support a model in which nSMase2 is expressed at the onset of neural crest EMT to produce ceramide and induce membrane curvature, thus increasing endocytosis of Wnt and BMP signaling complexes and activating pro-migratory gene expression. These results highlight the critical role of plasma membrane lipid metabolism in regulating transcriptional changes during developmental EMT programs.

## Introduction

Epithelial-to-mesenchymal transition (EMT) is a critical process in both development and metastasis during which epithelial cells gain migratory capacity to disperse throughout the body^1–3^. The transcriptional circuits that regulate these events have been at least partially elucidated and appear similar in both developmental and metastatic contexts^1–3^. Moreover, recent studies in cancer cells have begun to uncover significant changes in the plasma membrane lipid content during cancer cell EMT^4,5^. These observations raise the intriguing possibility that lipid metabolism functions to promote the EMT process by exploiting the structural and bioactive features of plasma membrane lipids. Accordingly, sphingolipid metabolism induces EMT in cancer cell lines by reducing ceramide content to suppress apoptosis and by producing the signaling lipids sphingosine-1-phosphate (S1P) and lysophosphatidic acid (LPA) to enhance proliferation and motility of metastatic cells^8–12^. Critical roles of bioactive lipids during development have also recently emerged. For example, signaling by S1P and LPA also regulate developmental cell migration ^13–15^, and the bioactive lipid ceramide induces neural tube closure defects when present at ectopic levels^16^. However, the functions of plasma membrane lipids during the developmental EMTs that precede migration remain less clear.

To determine how changes in plasma membrane lipid composition regulate developmental EMT *in vivo*, here we examined the avian neural crest, which undergoes a stereotypical EMT during development^3,17,18^. Specified premigratory neural crest cells exhibit epithelial characteristics and enter EMT under temporal control by a well-characterized gene regulatory network to adopt a migratory, mesenchymal physiology^17,19^, making it an excellent model to test how membrane lipid changes facilitate the transition to a mesenchymal fate. Evidence from the neural crest-derived pediatric cancer, neuroblastoma, suggests that lipid metabolism plays a critical role in neural crest biology^20,21^. In neuroblastoma, receptor tyrosine kinase activation is mediated by the organization of Src kinases into ordered plasma membrane microdomains^20^. Formation of ordered microdomains is largely driven by the thermodynamic features of membrane lipids^22^, thus highlighting the importance of understanding plasma membrane lipid metabolism during neural crest cell development.

Here we report that the sphingolipid metabolizing enzyme nSMase2 is necessary for neural crest EMT and migration. Mechanistically, our results suggest that nSMase2 activity enables endocytosis of cell signaling complexes from the plasma membrane, thereby potentiating Wnt and BMP signaling and activating the mesenchymal gene expression program critical for execution of the EMT to a migratory phenotype.

## Results

### Neural crest cells express Smpd3 during epithelial-to-mesenchymal transition and migration

We predicted that temporally regulated changes in lipid content during EMT would result from differential expression of lipid metabolic genes, we explored transcriptome resources comparing chicken cranial neural crest cells at premigratory (epithelial) and migratory (mesenchymal) stages^23^. We parsed these data for genes with Gene Ontology terms^24^ containing “lipid” and found six lipid-related genes significantly enriched in epithelial, premigratory neural crest, and nine significantly enriched in mesenchymal, migratory neural crest (Fig. 1a,b), consistent with a potential role for lipid modifications during neural crest EMT. Of particular interest, the gene *SMPD3*, which encodes the metabolic enzyme neutral sphingomyelinase 2 (nSMase2), showed the most significant enrichment at migratory stages (Fig. 1a). nSMase2 is a sphingolipid-metabolizing enzyme that functions at the plasma membrane to generate ceramide^6,7,25^. Since sphingolipid metabolism is tightly linked with EMT in cancer^4,10,26^, we delved further into the role of nSMase2 function during developmental neural crest EMT.

**Fig 1:**
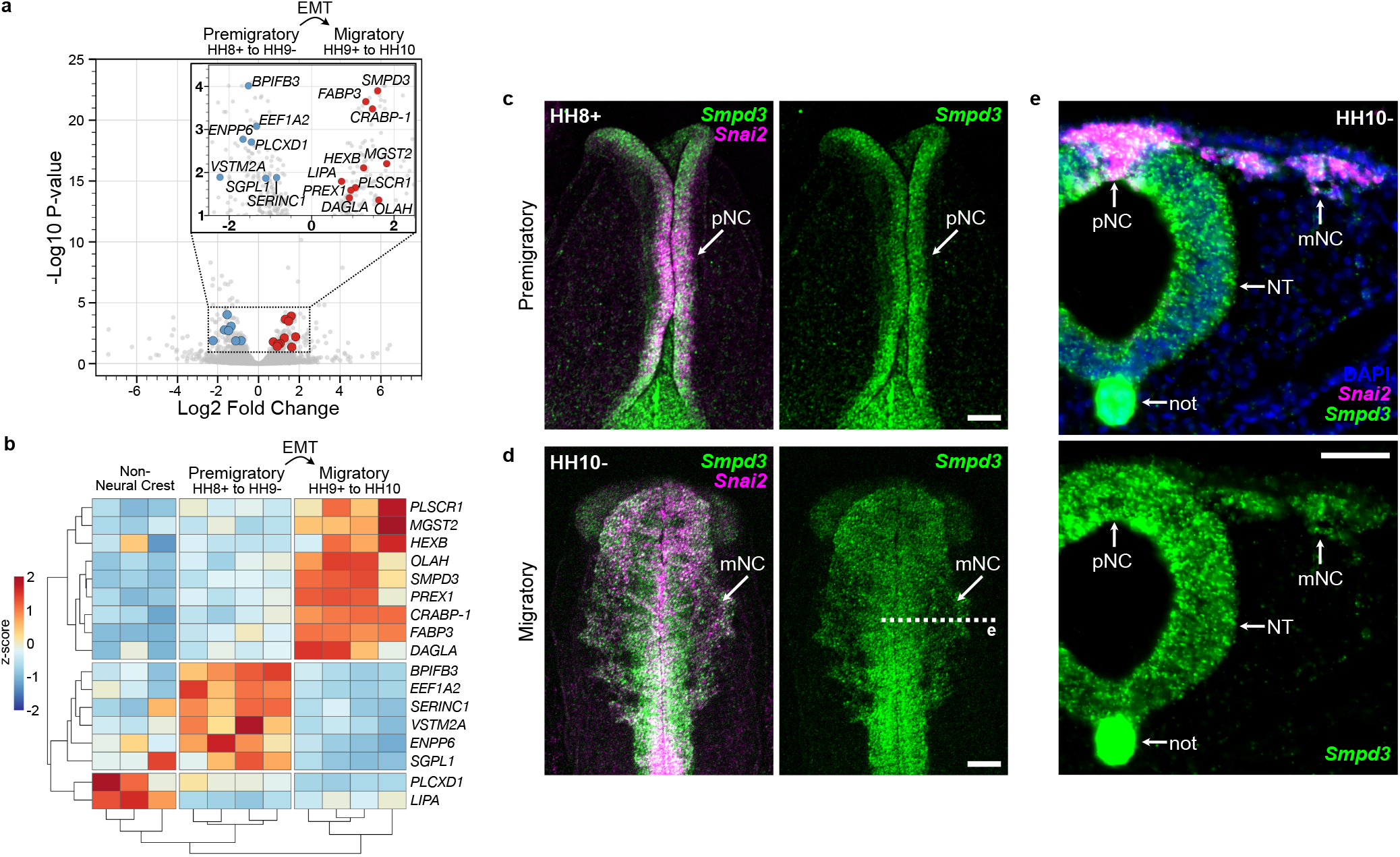
Identification of *Smpd3* enrichment during neural crest epithelial-to-mesenchymal transition. **a**, Volcano plot showing genes significantly enriched within premigratory (negative values) and migratory (positive values) neural crest cells. Lipid-related genes with significant differential gene expression are labeled (blue, premigratory enrichment; red, migratory enrichment). Data analysis was performed using previously published RNA seq reads^23^. EMT, epithelial-to-mesenchymal transition; HH, Hamburger-Hamilton stage. **b**, Heatmap displaying complete Euclidean clustering shows changes in gene expression levels for the lipid-related genes labeled in **a** between non-neural crest, premigratory, and migratory neural crest samples. Each column represents a single biological replicate. **c-d,** Hybridization chain reaction shows *Smpd3* and neural crest EMT marker *Snai2* expression at premigratory (**c**) and migratory (**d**) stages. pNC, premigratory neural crest; mNC, migratory neural crest. **e**, Transverse section through **d**(dashed line) shows strong *Smpd3* expression in the neural crest, as well as in the ventral neural tube (NT) and notochord (not). Scale bars represent 100 μm in **c**,**d** and 50 μm in **e**. See also Supplemental Fig. 1.

We first examined the spatiotemporal expression pattern of *Smpd3*, together with the neural crest marker *Snai2*, using Hybridization Chain Reaction (HCR), and found that *Smpd3* expression initiated prior to EMT in the specified neural crest cells at premigratory stages (HH8+) and remained highly expressed following EMT in neural crest cells at migratory stages (HH10−, Fig. 1c-e). We also observed strong expression of *Smpd3* in the more ventral neural tube and in the notochord (Fig. 1e). Chromogenic *in situ* hybridization confirmed these observations and revealed that *Smpd3* expression was not detectable at stages earlier than stage HH8 (Supplemental Fig. 1a-e), indicating that *Smpd3* expression is initiated just prior to neural crest EMT.

As a neutral sphingomyelinase enzyme, the *SMPD3* gene product, nSMase2, catalyzes the hydrolysis of sphingomyelin within the plasma membrane to produce ceramide^6,25^; thus, we asked if ceramide is produced in *Smpd3*-expressing neural crest cells. Immunolabeling of stage HH11 chicken embryos with antibodies against ceramide and the neural crest marker Pax7 revealed strong enrichment of ceramide on the apical surface of the neural tube (Supplemental Fig. 1f), correlating with the enriched *Smpd3* gene expression in this tissue (Fig. 1e). Importantly, ceramide labeling also appeared enriched on the surface of migrating neural crest cells (Supplemental Fig. 1f), indicating that ceramide is indeed produced in the cranial neural crest. Together these results raise the intriguing possibility that temporally regulated sphingolipid metabolism by nSMase2 may play a critical role during neural crest EMT.

### Ceramide production is required for cranial neural crest migration

The co-expression of *Smpd3* and ceramide during EMT stages led us to hypothesize that the *SMPD3* gene product, nSMase2, is required for ceramide production in the neural crest. To test this, we performed nSMase2 knockdown through bilateral electroporation of a non-binding control antisense morpholino (MO) on the left and an nSMase2-targetting MO on the right sides of gastrulating chicken embryos. Electroporated embryos were then allowed to develop to migratory stages, at which point neural crest cells were visualized (Fig. 2a). Compared to the contralateral control side, nSMase2 knockdown resulted in significantly decreased ceramide immunolabeling (Fig. 2b), demonstrating that nSMase2 is necessary for ceramide production in cranial neural crest cells.

**Fig. 2:**
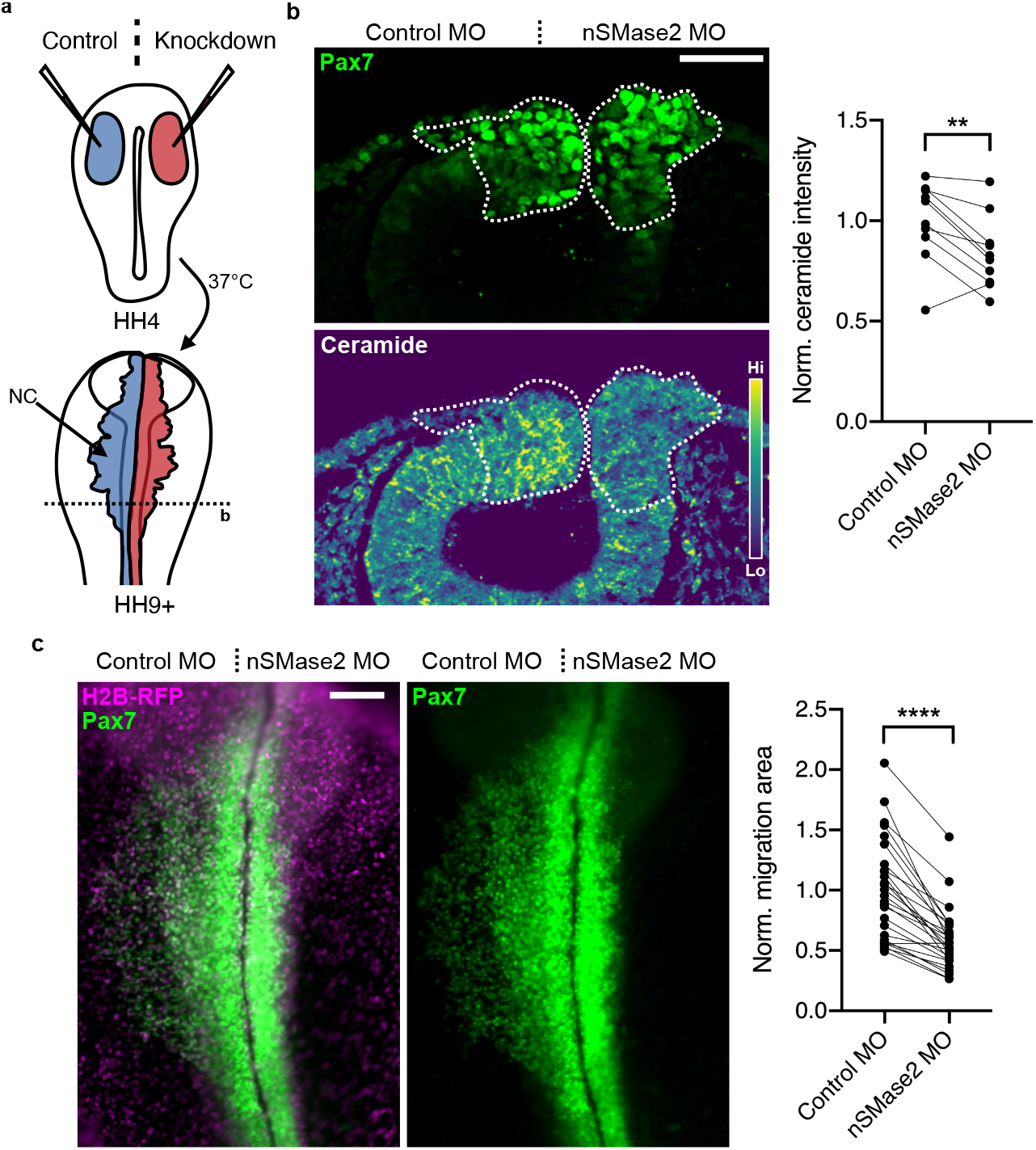
nSMase2-mediated ceramide production is required for neural crest EMT. **a**, Control (blue) and knockdown (red) reagents were delivered bilaterally into gastrulating chicken embryos (stage HH4), then embryos were incubated to neural crest migration stages for all *in vivo* electroporation experiments. Perturbations (right side) were then compared to the contralateral internal control side (left). **b**, Embryos were electroporated with a non-binding control morpholino (Control MO, left) and nSMase2-targeting morpholino (nSMase2 MO, right), then Pax7 and ceramide were immunolabeled in transverse sections through the cranial region at HH9. Parallel coordinate plot shows reduced ceramide immunoreactivity within the cranial neural crest region following nSMase2 knockdown (dashed lines in **b**). **c**, Control or nSMase2 MOs were electroporated together with H2B-RFP to display transfection efficiency. Pax7 immunolabeling shows failure of neural crest cells to migrate away from the midline following nSMase2 knockdown. Parallel coordinate plot displays normalized Pax7+ neural crest migration area, demonstrating significantly reduced neural crest migration following nSMase2 MO electroporation. Scale bars represent 50 μm (**b**) and 100 μm (**c**). \*\**p*<0.01, \*\*\*\**p*<0.0001, two-tailed paired *t*-test. Each point in **b** represents the mean of three sections from an individual embryo (n=10 embryos), and each point in **c** represents one embryo with lines connecting measurements from the same embryo (n=27 embryos).

Interestingly, Pax7 staining showed control neural crest cells successfully undergoing EMT and migrating away from the midline, while loss of nSMase2 on the contralateral side of the same embryos resulted in neural crest cells accumulating at the dorsal midline (Fig. 2b). This observation suggested that nSMase2-dependent ceramide production is required for neural crest migration. We next examined nSMase2 knockdown embryos in whole mount and quantitated the migration area of cranial neural crest cells on the control versus nSMase2 knockdown sides. While the control MO did not disrupt neural crest migration away from the midline, neural crest cells that received the nSMase2 MO showed a significant reduction in migration distance and these cells appeared to accumulate at the midline in the dorsal neural tube (Fig. 2c), consistent with our observations in transverse section. Importantly, markers of mitosis and apoptosis were unaffected by nSMase2 knockdown within the neural crest (Supplemental Fig. 2), and thus aberrant proliferation or survival does not explain the observed reduction in migration.

To determine if the effect of nSMase2 MO is specific to a loss of nSMase2, we performed a second independent method of nSMase2 knockdown using a CRISPR/Cas9-based approach^27^. Compared to a non-targeted gRNA, electroporation of Cas9 with nSMase2-targetting gRNA induced a significant reduction in *Smpd3* transcripts (Supplemental Fig. 3a), consistent with mutation-induced nonsense-mediated decay^27,28^. We then quantitated the migration area of cranial neural crest cells and observed a similar reduction in migration following CRISPR-mediated nSMase2 knockdown compared with nSMase2 MO (Supplemental Fig. 3b). Together these results demonstrate that nSMase2 function is critical for cranial neural crest migration and suggest that ceramide production by nSMase2 is required for EMT *in vivo*.

### nSMase2 is required to activate pro-migratory gene expression changes

The observed loss of cranial neural crest migration upon nSMase2 knockdown may result from a defect in activating migratory gene regulatory circuits or from disruption of the downstream cellular mechanisms underlying neural crest migration. To differentiate between these possibilities, we performed HCR to examine the expression of multiple markers that reflect neural crest specification (*Msx1*) and EMT progression (*Sox9* and *Snai2*)^17,19^. The activation of *Snai2* expression during EMT directly represses expression of the premigratory cadherin *CAD6B* in neural crest cells^29^, and thus we also examined expression of *Cad6b* as a premigratory marker in nSMase2 morphants. nSMase2-deficient neural crest displayed significant downregulation of *Msx1*, *Sox9*, and *Snai2* expression, and elevated expression of *Cad6b* transcripts (Fig. 3a, Supplemental Fig. 4). We next performed immunolabeling for Snail2 and Cad6B and, consistent with the HCR results, nSMase2 knockdown reduced Snail2 and elevated Cad6B protein levels in the cranial neural crest (Fig. 3b). Together these results indicate that nSMase2 is necessary to activate pro-EMT gene expression, most dramatically upregulating *Sox9* and *Snai2* expression. These observations suggest that nSMase2 activity promotes the transition from premigratory to migratory neural crest gene signatures during EMT, and the loss of migration observed in nSMase2-deficient neural crest reflects a failure to activate the mesenchymal gene expression program.

**Fig. 3:**
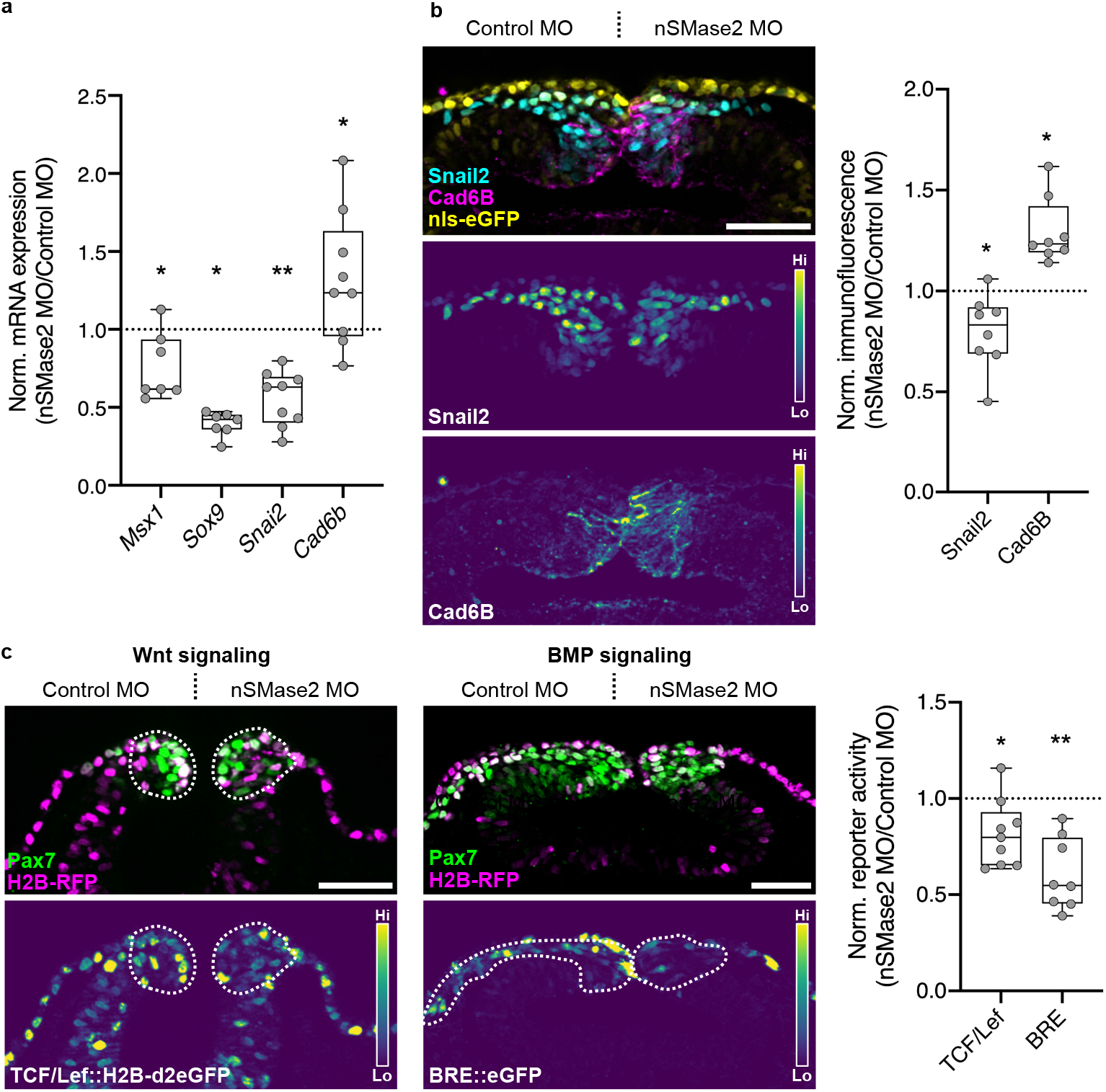
nSMase2 is required for the transcriptional activation of the EMT program and for Wnt and BMP signaling. **a**, Relative mRNA expression for neural crest markers in nSMase2 knockdown embryos at stage HH9− (representative images displayed in Supplemental Fig. 4, *Msx1* n=7 embryos, *Sox9* n=7 embryos, *Snai2* n=9 embryos, *Cad6b* n=7 embryos). **b**, Immunolabeling for Snai2 and Cad6B proteins in nSMase2 knockdown sections at HH9. Boxplot shows relative Snai2 and Cad6B protein levels as measured by fluorescence intensity (Snail2 n=8 embryos, Cad6B n=8 embryos). **c**, nSMase2 knockdown reagents were electroporated together with the Wnt-responsive TCF/Lef∷H2B- d2EGFP (left panel) or the BMP-responsive BRE∷eGFP (center panel) constructs and imaged in transverse sections at HH9− and HH10−, respectively. Relative eGFP fluorescence intensity within the Pax7+ neural crest domain shows a significant reduction in Wnt and BMP output in nSMase2-depleted neural crest cells (TCF/Lef n=9 embryos, BRE n=8 embryos). Scale bars represent 50 μm. \**p*<0.05, \*\**p*<0.01, two-tailed paired *t*-test. Each point in **a** represents one embryo, and in **b**,**c** represents the mean of three sections from individual embryos.

### Wnt and BMP signaling pathways require nSMase2 function

Activation of the EMT gene regulatory network in neural crest involves the canonical Wnt and BMP pathways^17,19,30^, and we hypothesized that sphingolipid metabolism at the plasma membrane may alter the activity of these pathways^7,31^. To test this, we turned to GFP reporter constructs which measure canonical Wnt (TCF/Lef∷H2B-d2eGFP) and BMP (BRE∷eGFP) pathway output in cranial neural crest cells^32–35^. We electroporated these reporters together with control or nSMase2 MOs, then compared GFP intensity within the cranial neural crest. At the onset of EMT at stage HH9−, we observed a significant reduction in Wnt/β-catenin signaling upon nSMase2 knockdown (Fig. 3c). That *SNAI2* is a direct target of Wnt signaling^36^ explains the observed loss of *Snai2* expression (Fig. 3a). Similarly, we observed a significant reduction in BMP signaling activity (Fig. 3c), consistent with the reduced expression of *Msx1*(Fig. 3a), a downstream target of BMP signaling^37^. Together these results demonstrate that nSMase2 function is required for signaling cascades, including the Wnt and BMP pathways, originating from the plasma membrane.

### nSMase2 expression is sufficient to induce endocytosis

We next sought to mechanistically test the role of sphingolipid metabolism at the plasma membrane in the activation of developmental signaling. Wnt and BMP signaling components collect within sphingolipid- and cholesterol-rich membrane microdomains, such as caveolae^22,38,39^. Ceramide production within such microdomains is sufficient to induce membrane curvature and internalization^40–43^, thus resulting in the internalization of microdomain-localized proteins. Importantly, endocytosis of Wnt and BMP/TGF-β complexes into signalosomes facilitates the most potent signaling output^39,44–47^. This raised the possibility that nSMase2 function promotes endocytosis of membrane microdomains, thus explaining the reduction in Wnt and BMP signaling following nSMase2 knockdown.

We tested this hypothesis by assaying endocytosis in the epithelial U2OS cell line that bears morphological similarity to premigratory neural crest. To this end, we employed fluorescently labeled Transferrin (Tf-633) as a readout for receptor-mediated endocytosis^48^. Cells were transfected with control or perturbation constructs, then exposed to Tf-633 for 30 minutes in the culture media before fixation, imaging, and quantitation of Tf-633-positive endocytic puncta. Compared with an RFP-only transfection control, transfection of a dominant-negative Dynamin 1 mutant (Dyn1(K44A))^49,50^ resulted in reduced Tf-633 internalization (Fig. 4a), as predicted. Next, we found that nSMase2 overexpression was sufficient to increase Tf-633 internalization (Fig. 4a), consistent with our hypothesis that nSMase2 expression in the premigratory neural crest activates endocytosis. Finally, we generated a mutant of nSMase2 (nSMase2(N130A)) lacking catalytic function ^51^, and found that without its catalytic activity, nSMase2(N130A) overexpression had no effect on endocytosis (Fig. 4a). These results support our model in which nSMase2 expression in epithelial cells, similar to premigratory neural crest cells, activates endocytosis through the conversion of sphingomyelin into ceramide and results in active cell signaling.

**Fig. 4:**
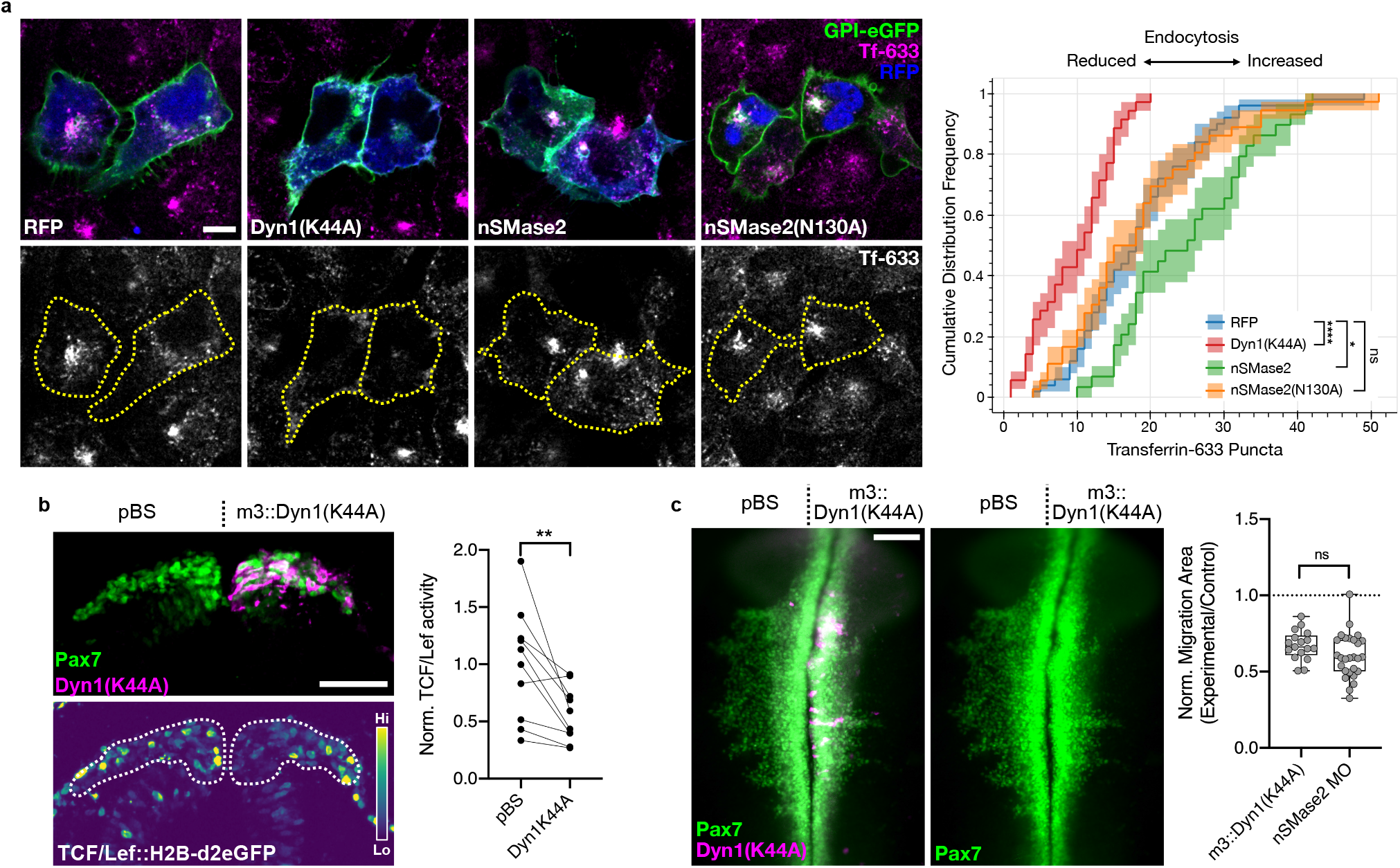
nSMase2 is sufficient to induce endocytosis, and endocytosis is necessary for neural crest EMT. **a**, Epithelial cells (U2OS) were transfected with GPI-eGFP to label the plasma membrane and the indicated overexpression constructs. After 24 hours, transfected cells were incubated with Transferrin-Alexa-633-conjugate (Tf-633) in the culture media for 30 minutes to allow for receptor-mediated endocytosis. Cells were then fixed and visualized by confocal microscopy. Scale bars represent 10 μm. Cumulative distribution frequency (CDF) plot displaying the number of fluorescent Tf-633 puncta within individual transfected cells. CDF curves shifted to the left are consistent with reduced endocytosis, while curves shifted to the right are indicative of elevated endocytic activity (n=50 RFP, 35 Dyn1(K44A), 29 nSMase2, and 36 nSMase2(N130A) cells). ns not significant, \**p*<0.05, \*\*\*\**p*<0.0001; Kruskal-Wallis test (*p*=2.69e-9) with Dunn’s posthoc analysis. **b**, The Wnt-responsive TCF/Lef∷H2B-d2eGFP reporter was coelectroporated with the dominant negative Dyn1(K44A) under control of the neural crest-specific *FoxD3* NC1.1m3 enhancer (m3∷Dyn1(K44A)). Embryos were collected and examined in transverse section at HH9+. Normalized eGFP fluorescence quantitation shows reduced Wnt signaling following Dyn1(K44A) expression compared to a non-expressing pBluescript (pBS) control (n=10 sections from 3 embryos). \*\**p*<0.01; two-tailed paired *t*-test. Scale bar represents 50 μm. **c**, Embryos were electroporated with m3∷Dyn1(K44A) and examined in whole mount at HH9+ for neural crest migration by Pax7 immunolabeling. Neural crest migration was quantitated from endocytosis-inhibited embryos, revealing a reduction in migration area. Comparing relative migration area between m3∷Dyn1(K44A) embryos with nSMase2 knockdown embryos shows that endocytosis inhibition phenocopies nSMase2 knockdown (n=17 m3:Dyn1(K44A) and 27 nSMase2 MO embryos). ns not significant; two-tailed unpaired *t*-test. Each point represents one embryo. Scale bar represents 100 μm. nSMase2 MO data is reproduced here from Fig. 2c.

### Inhibition of endocytosis phenocopies loss of nSMase2 during neural crest EMT

To test this model *in vivo*, we asked if endocytosis is required for neural crest EMT. If so, inhibition of endocytosis within the neural crest during EMT should reproduce the phenotypes observed following nSMase2 knockdown. To this end, we inhibited endocytosis specifically within cranial neural crest cells by expressing the dominant negative Dyn1(K44A) mutant under control of the neural crest-specific *FoxD3* NC1.1m3 enhancer^52^ (m3∷Dyn1(K44A)). Following electroporation of m3∷Dyn1(K44A), we observed reduced Wnt/β-catenin signaling at EMT stages within the cranial neural crest cells (Fig. 4b). We then examined neural crest migration following m3∷Dyn1(K44A) electroporation and found that endocytosis was required during EMT for cranial neural crest migration (Fig. 4c). These data together indicate that endocytosis is required during neural crest EMT for Wnt/β-catenin signaling and for migration, thus phenocopying nSMase2 knockdown and suggesting that endocytosis acts downstream of nSMase2 function during neural crest EMT.

## Discussion

Here we show that premigratory neural crest cells initiate expression of nSMase2 to convert plasma membrane sphingomyelin into ceramide in preparation for EMT. Ceramide production has strong effects on lateral membrane organization by forming microdomains^22,53–55^ that drive receptor clustering and plasma membrane curvature^40,55^. We find that the production of ceramide is required for Wnt and BMP signaling during neural crest EMT, likely by promoting endocytosis of activated signaling complexes^38,39,44–47,56^. Accordingly, we find that nSMase2 is sufficient to induce endocytosis, and inhibition of endocytosis mimics the effects of nSMase2 knockdown in cranial neural crest EMT. Thus, our results suggest that temporally regulated ceramide production by nSMase2 mechanistically activates endocytosis to facilitate neural crest EMT and migration (Fig. 5). The results reveal that changes in lipid metabolism are a critical driver of developmental cell signaling that is required for the transition to a migratory phenotype.

**Fig. 5:**
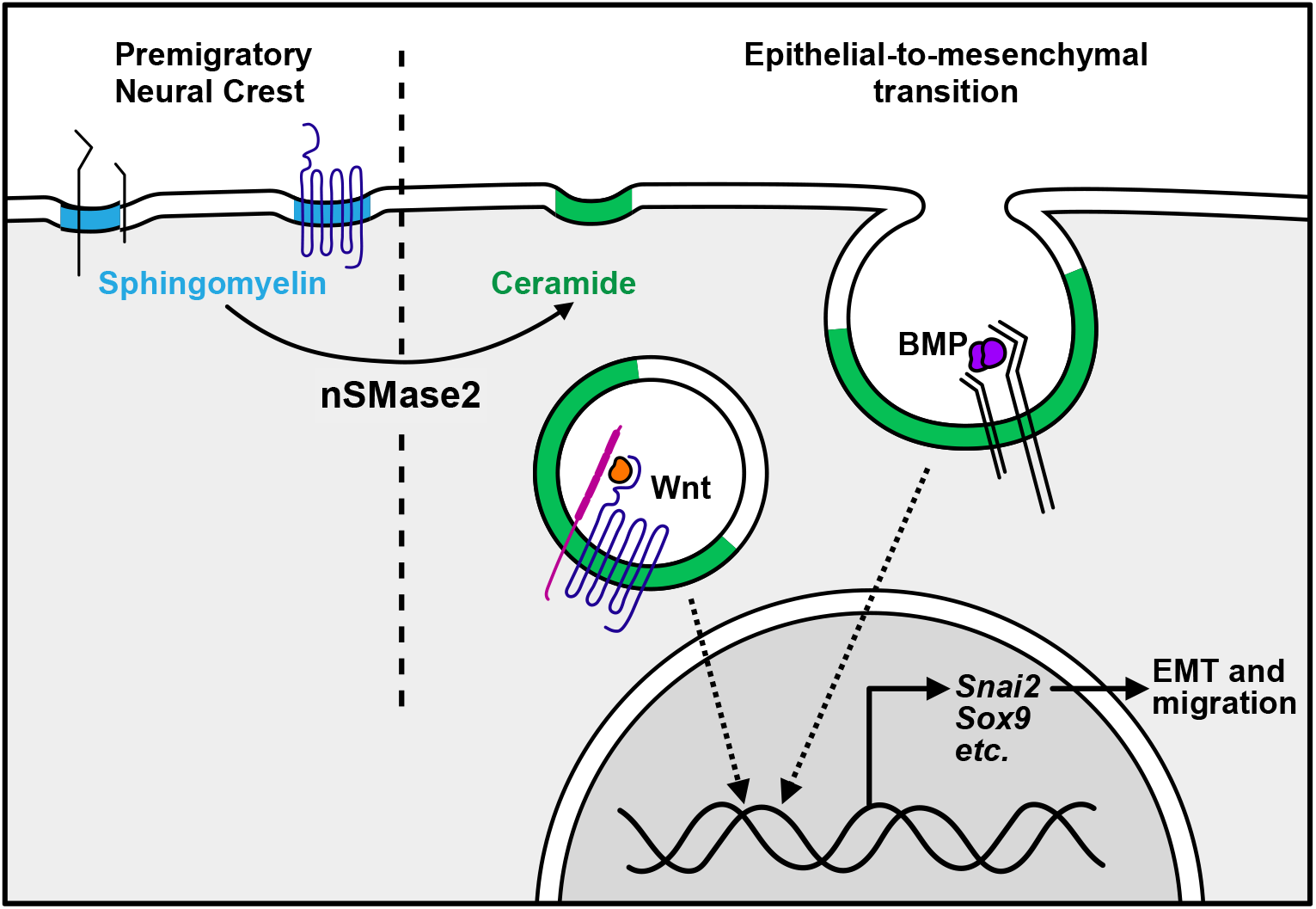
nSMase2 regulates endocytosis to enable neural crest EMT. Premigratory neural crest cells have sphingomyelin-rich membrane microdomains. At the onset of EMT, nSMase2 expression converts sphingomyelin into ceramide to promote membrane curvature. This facilitates the endocytosis of activated signaling complexes (Wnt and BMP). Internalization of these complexes enables activation of pro-EMT transcriptional targets (including *Snai2* and *Sox9*) which promote neural crest EMT and migration.

In cancer cell lines, ceramide production shows pro-apoptotic effects, thereby inhibiting EMT; instead, EMT and motility are enabled by the downregulation of ceramide or the upregulation of sphingomyelin production^8–10^. These observations contrast with our findings during developmental EMT, where ceramide production instead activates EMT gene expression (Fig. 3) and motility (Fig. 2c) without affecting proliferation or apoptosis (Supplemental Fig. 2). Our model suggests that the removal of sphingomyelin is necessary for mesenchymalization during development. Consistent with this, in collecting duct cells sphingomyelin synthesis is required to maintain epithelial properties^57^. Together, these observations suggest that sphingomyelin and ceramide have opposite effects in developmental versus metastatic EMT progression. It will be of great interest to determine the context-specific mechanisms that regulate this differential response.

The observation that nSMase2 function increases Transferrin endocytosis (Fig. 4a) is very intriguing given the lateral organization of Transferrin receptors on the plasma membrane. Transferrin binds preferentially within disordered membrane microdomains^58^, whereas sphingomyelin and ceramide are enriched within more ordered microdomains^53^. How sphingomyelin metabolism in ordered microdomains causes endocytosis of disordered microdomain-localized proteins remains an unanswered question. There is evidence that SMase function enables endocytosis of ordered caveolar membrane microdomains. Sphingomyelin within caveolae stabilizes plasma membrane invaginations, and treatment with SMase decreases sphingomyelin content and reduces caveolae stability^43^. Since activated Wnt and BMP receptors localize to caveolae^22,38,39^, this supports the notion that nSMase2-dependent conversion of sphingomyelin into ceramide in neural crest cells enables internalization of Wnt and BMP receptor complexes within caveolae. Together, these findings highlight how dynamic changes in lipid metabolism alter plasma membrane function to enable endocytosis and have broad reaching effects on critical developmental processes such as EMT and cell migration.

## Acknowledgements

We would like to thank Megan Martik, Erdinç Sezgin, Justin Bois, and Steven Wilbert for valuable discussion on experiment design and analysis, and Alexis Camacho-Avila and Gabriel da Silva Pescador for technical support. We thank Catherine Berlot, Anna-Katerina Hadjantonakis, Randall Moon, and Elisa Martí for sharing reagents. Confocal imaging was supported by the Caltech Beckman Institute and the Arnold and Mabel Beckman Foundation to the Biological Imaging Facility. Funding for this work comes from the National Institutes of Health grants K99DE029240 to M.L.P., K99DE028592 to E.J.H, R01DE027538 and R01DE027568 to M.E.B, and from the Caltech Summer Undergraduate Research Fellowship (SURF) to C.J.A.

## Author Contributions

Conceptualization: M.L.P. and M.E.B.

Experiment design: M.L.P., E.J.H., and M.E.B.

Experimentation: M.L.P., E.J.H., and C.J.A.

Data analysis: M.L.P. and C.J.A.

Data interpretation: M.L.P., E.J.H., and M.E.B.

Manuscript preparation: M.L.P. and M.E.B.

Manuscript editing: E.J.H.

## Data Availability

All data that support the findings of this study are available from the corresponding author upon reasonable request. The *Smpd3* mRNA sequence has been submitted to GenBank (Accession # Pending).

## Code Availability

All data analysis code used in this study are available from the corresponding author upon reasonable request.

## Materials and Methods

### RNA Seq Analysis

Bulk RNA Seq reads from FACS-isolated *FoxD3*-positive premigratory (HH8+ to HH9−) and migratory (HH9+ to HH10) chicken cranial neural crest cells, and *FoxD3*-negative non-neural crest cells, were downloaded from GEO (BioProject PRJNA497574)^23^. Reads were trimmed using Cutadapt^59^ and aligned to the chicken genome (GRCg6a) using BowTie2^60^. Transcripts were then counted using featureCounts^61^ and differential expression analysis between premigratory and migratory gene expression was carried out using DESeq2^62^. Volcano plot analysis (Fig. 1a) highlights genes with lipid-related GO terms^24^ with an adjusted p-value < 0.05, base mean value > 45, and log2-fold change value > 0.7. Complete Euclidean distance clustering was performed using pheatmap^63^ (Fig. 1b), and displays each premigratory, migratory, and non-neural crest cell biological replicate.

### Embryos and perturbations

Fertilized chicken eggs were obtained from commercial sources (Sunstate Ranch, Sylmar, CA and AA Lab Eggs, Westminster, CA). *Ex ovo* electroporations were performed using five pulses of 5.6 V for 50 ms at 100 ms intervals and cultured in albumin with 1% penicillin/streptomycin, then incubated to the desired HH stage^64,65^. For transverse sections, fixed embryos were incubated in 5% sucrose for 30 minutes at room temperature, 15% sucrose overnight at 4°C, 7.5% gelatin at 39°C overnight, flash-frozen in liquid nitrogen, and cryosectioned at a thickness of 18 μm. All reagents electroporated either expressed a fluorescently-tagged protein or were co-electroporated with nuclear GFP- or RFP-expressing constructs^66,67^; fluorescent protein expression was used to screen for electroporation efficiency and only embryos with high efficiency were included in analysis. Morpholinos were electroporated at 0.8 mM and synthesized with the following sequences (Gene Tools): control MO (5’-CCTCTTACCTCAGTTACAATTTATA-3’), translation-blocking nSMase2 MO (5’-GGTGTCACTGTGTCAAGCATCCATA-3’). pTK-*FoxD3*-NC1.1m3∷Dyn1(K44A) over-expressing constructs were electroporated at 1.5 μg/μl. CRISPR-mediated knockdown was performed using a plasmid-based strategy, and a nonbinding control gRNA (5’-GCACTGCTACGATCTACACC-3’), and a nSMase2-targeting gRNA designed within the first exon of the *SMPD3* gene locus (5’-GCAATCTGCGCAGCCCGAGA-3’), each at 1.5 μg/μl concentrations, were co-expressed with a Cas9-expressing plasmid (1.5 μg/μl) as previously described^27^.

### Hybridization chain reaction and in situ hybridization

Reagents for third-generation hybridization chain reaction were purchased from Molecular Technologies, and hybridization experiments were performed following manufacturer’s instructions^68^. Probe sets were designed against *Smpd3*(B1 initiator), *Snai2*(B4), *Cad6b*(B5), *Sox9*(B5), *Msx1*(B4), and *Tfap2b*(B7), and detected using appropriate amplifier hairpins labeled with Alexa488, Alexa546, and Alexa647. For chromogenic *in situ* hybridization, a 970 bp fragment of the chicken *Smpd3* transcript, corresponding to 76 bp of the 5’ untranslated region and first 894 bp of open reading frame was amplified by PCR from stage HH9 cDNA using primers ‘nSMase2 FWD −76’ and ‘nSMase2 REV 894 T7’ (Supplemental Table 1). RNA probes were synthesized and labeled with DIG (Roche) using the T7 RNA polymerase (Promega) and purified with Illustra Probe-Quant G-50 Micro Columns (GE Healthcare). Hybridization and probe detection were carried out as previously described^35^.

### Immunohistochemistry

For immunofluorescence analysis, embryos were fixed for 20 minutes at room temperature in 4% PFA in phosphate buffer, and all subsequent washes and incubations were performed in TBST + Ca^2+^ (50 mM Tris-HCl, 150 mM NaCl, 1 mM CaCl_2_, 0.5% Triton X-100). Blocking was performed in 10% donkey serum for 2 hours at room temperature, and primary and secondary antibody incubations were carried out in 10% donkey serum for 2 nights at 4°C. Primary antibodies employed in this study include mouse IgG1 anti-Pax7 (1:10; DSHB #PAX7), mouse IgM anti-HNK-1 (1:5; DSHB #3H5), mouse IgG1 anti-Cad6B (1:5; DSHB #CCD6B-1), mouse IgM anti-ceramide (1:200; Sigma #MID15B4), rabbit anti-phosphohistone H3 (1:500; EMD Millipore #06-570), rabbit anti-cleaved caspase 3 (1:300; R&D Systems #AF835), rabbit anti-Snai2 (1:500; Cell Signaling Technologies #9585), goat anti-GFP (1:500; Rockland #600-101-215M), and rabbit anti-RFP (1:500; MBL #PM005). Alexa Fluor 350-, 488-, 568-, 633-, or 647-conjugated donkey secondary antibodies (1:500; Molecular Probes) were used to detect primary antibodies.

### Construct design and cloning

Cloning was performed by a combination of PCR reactions with high-fidelity DNA polymerase AccuPrime (ThermoFisher), and restriction digestion and ligation (enzymes from NEB). A 5’ Kozak consensus sequence was introduced to promote efficient translation of each overexpression construct, and all constructs were sequence verified before use. All primer sequences are presented in Supplemental Table 1. CMV∷Dyn1(K44A)-mRFP (Addgene plasmid #55795) ^49^, pCAG∷GPI-eGFP (Addgene plasmid #32601)^69^, TCF/Lef∷GFP (Addgene plasmid #32610)^32^, M38-TOP∷d2EGFP (Addgene plasmid #17114) and BRE∷GFP^34^ constructs were generous gifts from Catherine Berlot, Anna-Katerina Hadjantonakis, Randall Moon, and Elisa Martí.

The nSMase2 open reading frame was amplified from cDNA using ‘nSMase2 ATG XhoI’ and ‘nSMase2 stop ClaI’ primers and ligated into pCI∷H2B-RFP^66^ between the XhoI and ClaI sites to produce pCI∷nSMase2. The nSMase2(N130A) mutation was inserted by amplifying the 5’ and 3’ fragments of nSMase2 from cDNA using ‘nSMase2 FWD −76’ and ‘nSMase2 N130A REV’, and ‘nSMase2 N130A FWD’ and ‘nSMase2 2462 REV’, respectively. These PCR fragments were then fused by amplification with ‘nSMase2 ATG XhoI’ and ‘nSMase2 stop ClaI’ and ligated into pCI∷H2B-RFP between the XhoI and ClaI sites, producing pCI∷nSMase2(N130A). pCAG∷nSMase2-RFP was produced by amplifying the nSMase2 open reading frame without the stop codon from pCI∷nSMase2 using ‘nSMase2 ATG SphI’ and ‘nSMase2 nostop ClaI’ primers, and amplifying RFP from pCI∷nSMase2 using ‘RFP ATG ClaI’ and ‘RFP stop NotI’ primers. These fragments were digested and ligated together between the SphI and NotI sites of pCI∷H2B-RFP. pCAG∷2a-RFP was produced to introduce a self-cleaving 2a peptide sequence immediately upstream of RFP. The RFP sequence was amplified from pCI∷H2B-RFP, and the 2a sequence was synthetically added by sequential amplification with ‘2a-RFP FWD 1’, then ‘2a-RFP FWD 2’, then ‘2a-RFP FWD 3 ClaI’, each with ‘RFP stop NotI’. The resulting 2a-RFP sequence was then ligated into pCI∷H2B-RFP between the ClaI and NotI cut sites. pTK-*FoxD3*-NC1.1m3∷Dyn1(K44A)-RFP was produced by digesting pTK-*FoxD3*-NC1.1m3∷GFP^52^ with NheI and XbaI. CMV∷Dyn1(K44A)-RFP was grown in One Shot INV110 *dam- E. coli*(ThermoFisher) to prevent XbaI methylation and was similarly digested with NheI and XbaI. The resulting products were ligated together and sequenced to confirm orientation. Finally, GFP was replaced in TCF/Lef∷H2B-GFP^32^ with d2EGFP from M38-TOP∷d2EGFP by digestion and ligation between the AgeI and NotI cut sites.

### Tissue culture, transfection, and endocytosis assays

Human osteosarcoma U2OS cells (ATCC #HTB-96) were cultured at 37°C in 50% CO_2_ in McCoy’s 5A Modified Media (ThermoFisher) supplemented with 10% fetal bovine serum (Gibco) and 1% penicillin/streptomycin (Corning). Cells were seeded for transfection in 8-well glass bottom chamber slides (ibidi # 80827) and transfected using Lipofectamine 3000 (Invitrogen) at 80% confluency then incubated for 24 hours. Culture media was replaced with 25 μg/mL Transferrin-Alexa-633-conjugate (ThermoFisher #T23362) in serum-free DMEM (ThermoFisher) and incubated at 37°C for 30 minutes. Cells were then washed 2x with DPBS (ThermoFisher) and fixed with 4% PFA in PBS for 10 minutes at room temperature before imaging.

### Microscopy and image analysis

Whole mount and transverse section imaging was performed using a Zeiss Imager.M2 with an ApoTome.2 module or with a Zeiss LSM 880 upright confocal microscope. All transverse section images, as well as whole mount embryo images (Fig. 1c,d, Supplemental Fig. 3a, and Supplemental Fig. 4) display maximum intensity projections of Z-stacks. Whole mount embryo images (Fig. 2c, Fig. 4c, and Supplemental Fig. 3b) display wide-field views. Cell culture experiments were imaged using a Zeiss LSM 800 inverted confocal microscope and displayed as single focal plane images. Images were analyzed and prepared for display using Fiji^70^. Regions of interest were manually drawn to measure fluorescence intensity and neural crest migration area, and to define boundaries for particle counting. For Tf-633 particle counting, images were subjected to median filtering, then thresholding using the Triangle method in the Auto Threshold tool. Analyze Particles was then used to count puncta with areas between 0.25 and 15.0 μm^2^. For PH3-positive cell counting, median filtered images were thresholded using the Bernsen method with the Auto Local Threshold tool and the Analyze Particles feature then was used to quantitate particles larger than 20.0 μm^2^. For cleaved-Casp3 quantitation, median filtered images were subjected to uniform manual thresholding and Analyze Particles was used to count puncta larger than 3 μm^2^.

### Statistical analysis

Parallel coordinate plots (Fig. 2b,c, Fig. 4b, and Supplemental Fig. 3b) show the control and experimental measurements, normalized to the mean of the control group, with solid lines connecting values from the same embryo. Box plots (Fig. 3, Fig. 4c, and Supplemental Fig. 3a) display the ratio of experimental divided by corresponding control measurements, with the following box plot elements: center line, median; box limits upper and lower quartiles; whiskers, minimum and maximum values; points, individual embryo measurements. Statistical analyses were performed using GraphPad Prism 8 software. All *t*-tests were two-tailed and paired, with the exception of one two-tailed, unpaired *t*-test (Fig. 4c). All embryo analyses show pooled results from at least three independent experiments, and all *t*-tests passed power analysis in G*Power 3.1^71^ at a power cutoff of 0.80. Cumulative distribution frequency analysis for Tf-633 endocytosis shows pooled results from two independent experiments and was performed in Python using iqplot^72^. Kruskal-Wallis test with Dunn’s posthoc analysis was used for statistical analysis of cumulative distribution frequencies.

---

**Supplemental Fig. 1:**
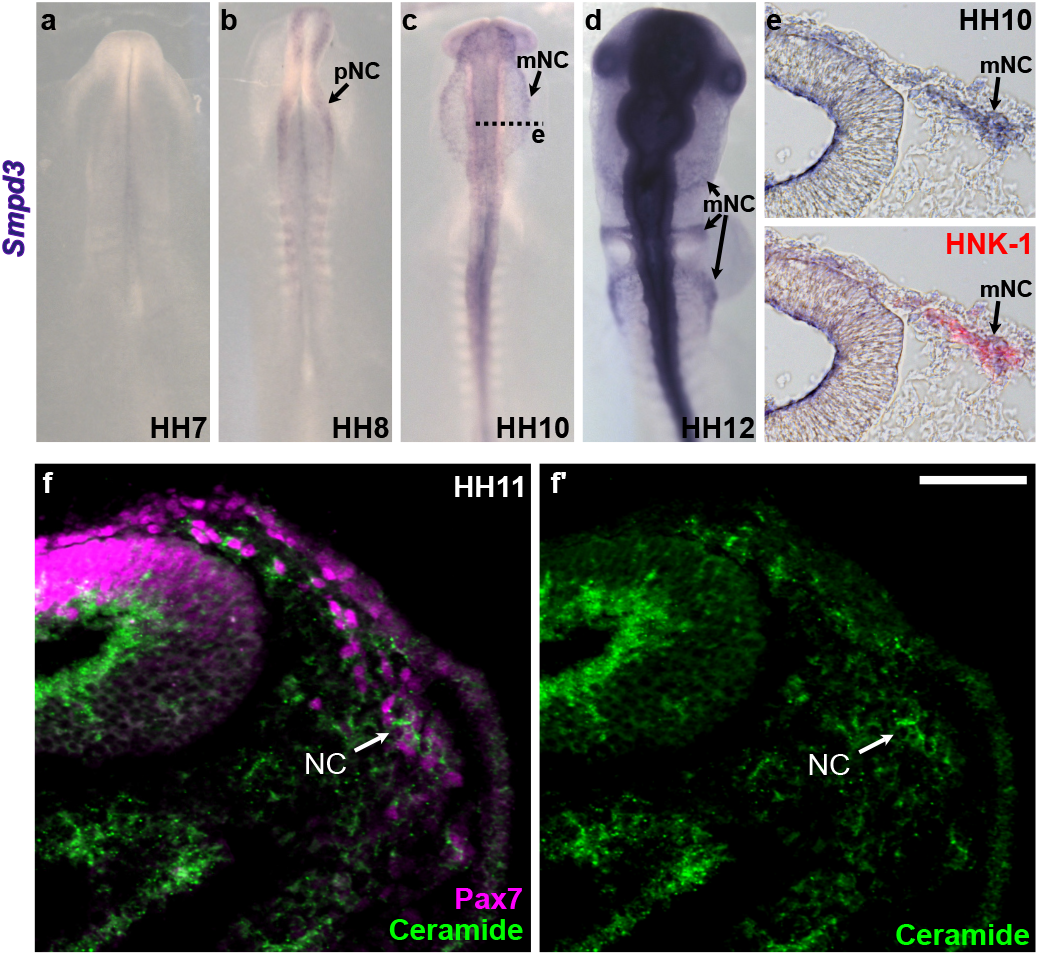
*Smpd3* gene expression is detected in premigratory and migratory cranial neural crest cells. *In situ* hybridization shows that *Smpd3* expression is absent at early stages (**a**), initiates in specified, premigratory neural crest cells (**b**), and is maintained during migration (**c**,**d**). **e**, Transverse section of embryos at stage HH10 (**c**, dashed line) shows *Smpd3* expression in migrating cranial neural crest cells, immunolabeled by the surface antigen HNK-1. HH, Hamburger Hamilton stage; pNC, premigratory neural crest; mNC, migratory neural crest. See also Fig. 1. **f**, Immunolabeling reveals enrichment of the lipid ceramide in Pax7-labeled migrating neural crest cells at stage HH11, as well as along the apical surface of the neural tube.

**Supplemental Fig. 2:**
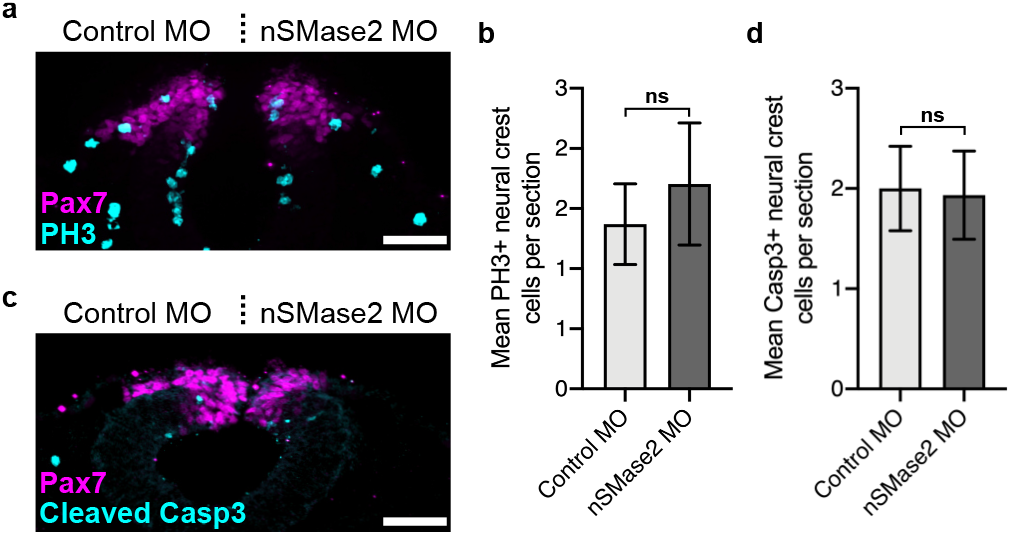
nSMase2 knockdown does not affect neural crest proliferation or survival. Gastrulating chick embryos were electroporated with a non-binding control morpholino (Control MO, left) and nSMase2-targeting morpholino (nSMase2 MO, right), and where subsequently immunostained for the neural crest marker Pax7 and the mitotic marker phospho-histone H3 (PH3, **a**), or for the apoptosis indicator, activated caspase 3 (cleaved Casp3, **c**) in section at the HH9+. Quantitation of PH3+ (n=27 sections from 9 embryos, **b**) and Casp3+ (n=15 sections from 5 embryos, **d**) cells show no significant change between nSMase2 knockdown and the contralateral control. Displayed is the mean with error bars reflecting sem. Scale bars represent 50 μm. ns, non-significant; two-tailed paired *t*-test.

**Supplemental Fig. 3:**
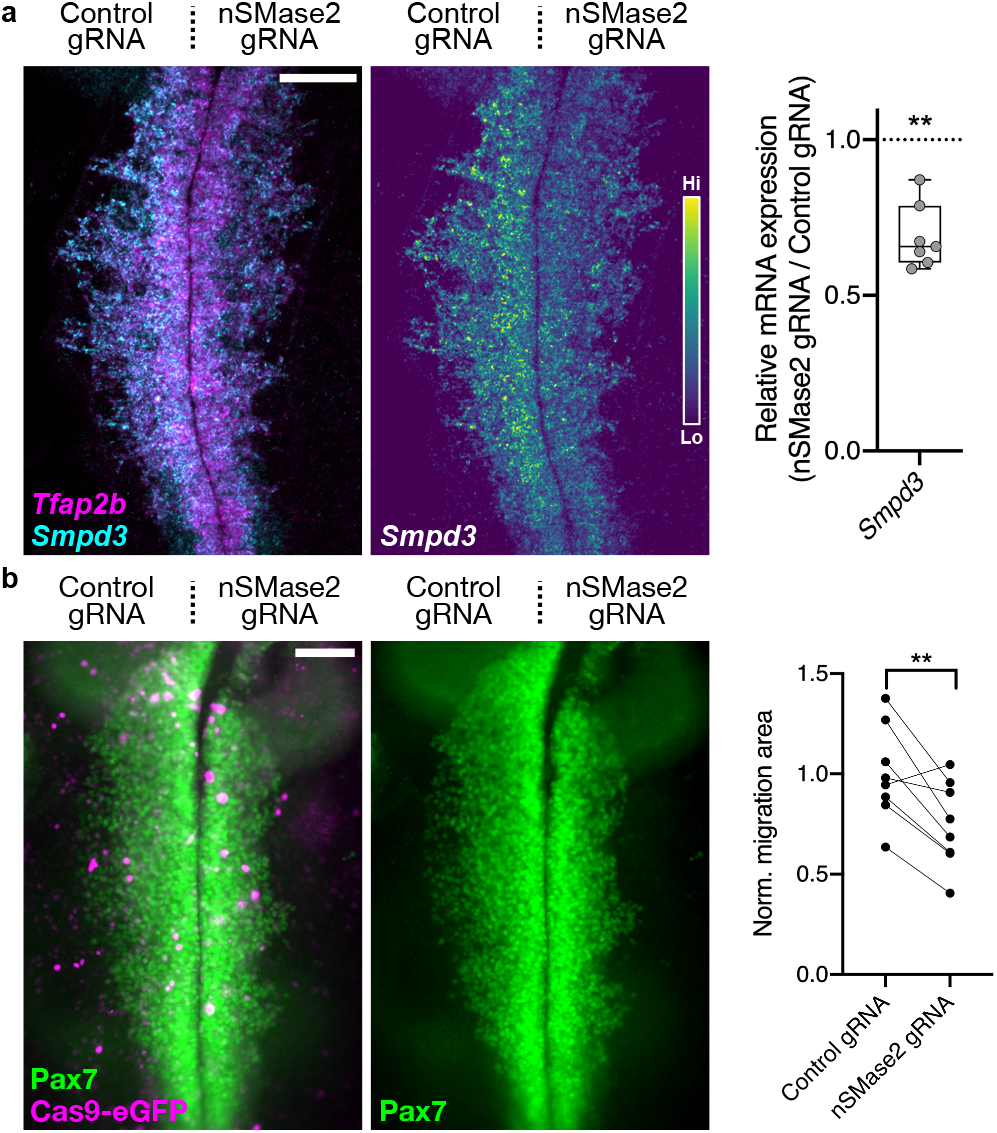
CRISPR-mediated nSMase2 knockout phenocopies nSMase2 MO. Embryos were electroporated with a Cas9-eGFP-expressing construct, together with U6.3-driven control- or nSMase2-targeting gRNAs. **a**, At HH9, embryos were fixed and processed for HCR to detect expression of *Smpd3* transcripts and the neural crest marker *Tfap2b*. Box plot shows relative fluorescent intensity shows reduced *Smpd3* expression following nSMase2 gRNA electroporation compared with control gRNA (n=7 embryos). **b**, Pax7 staining in nSMase2 CRISPR-electroporated embryos reveals a significant reduction in cranial neural crest migration area (n=8 embryos). Scale bars represent 100 μm. \*\**p*<0.01, two-tailed paired *t*-test. See also Fig. 2.

**Supplemental Fig. 4:**
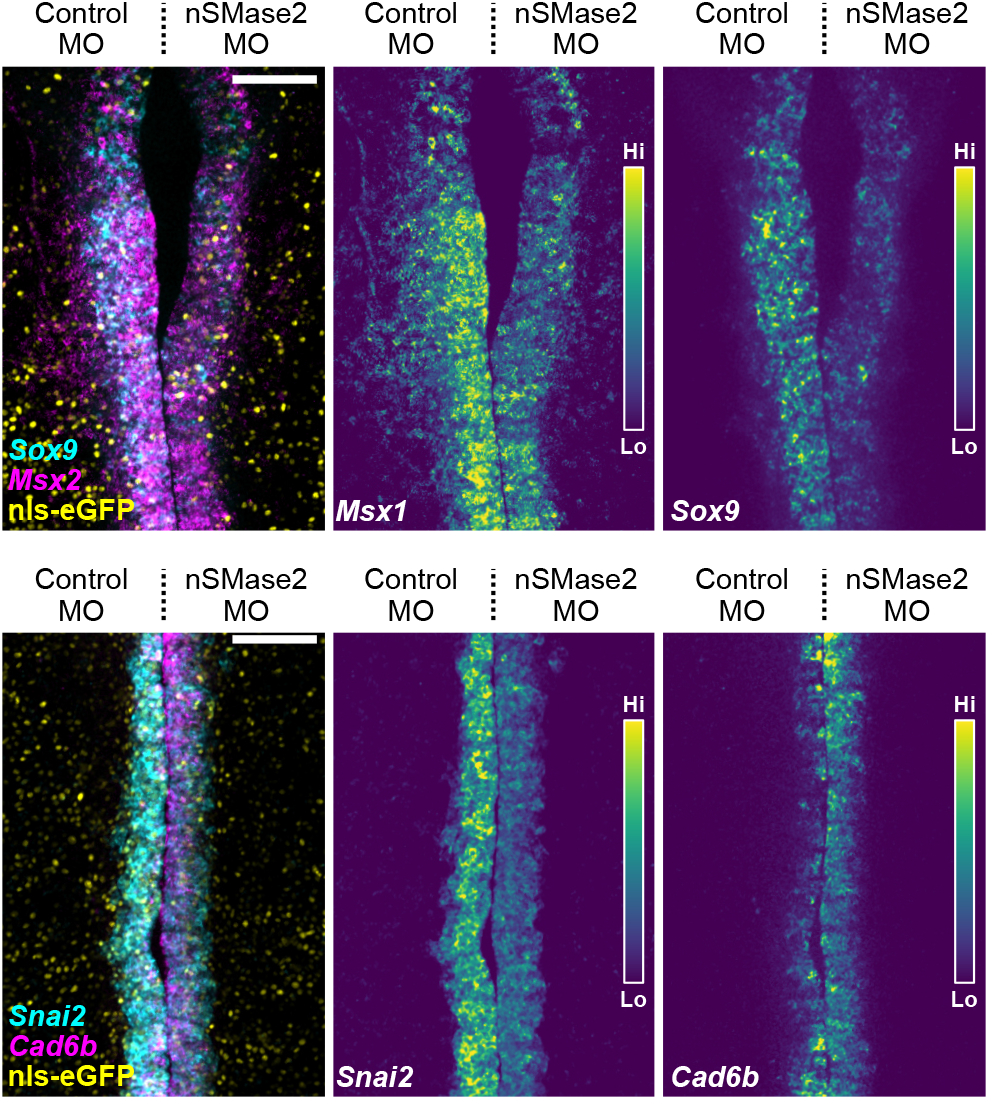
nSMase2 knockdown disrupts activation of the EMT transcriptional program. Representative images of nSMase2 knockdown embryos coelectroporated with nls-eGFP to label transfected cells, followed by HCR analysis for expression of neural crest markers *Msx1*, *Sox9*, *Sna2*, and *Cad6b* at stage HH9−. Scale bars represent 100 μm. See quantitation in Fig. 3a.

**Supplemental Table 1.**
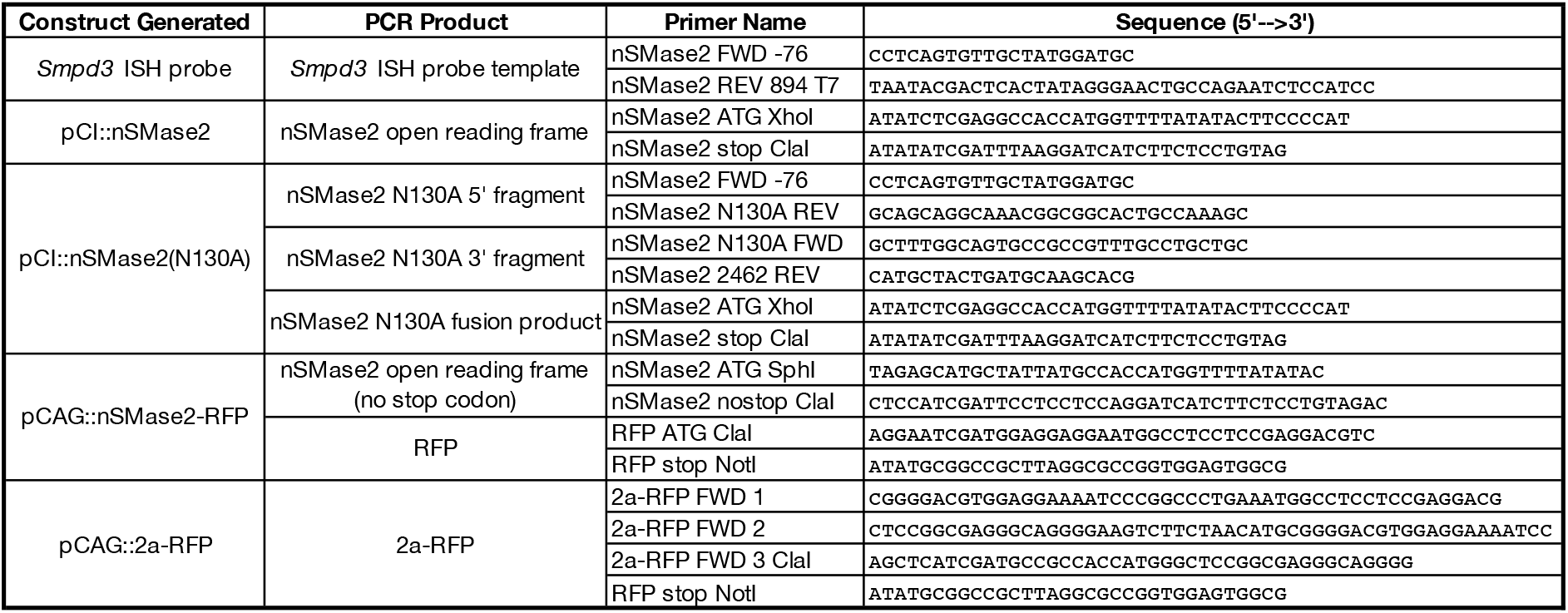
Primer sequences employed in this study.

